# Rapid functionalisation and detection of viruses via a novel Ca^2+^-mediated virus-DNA interaction

**DOI:** 10.1101/629303

**Authors:** Nicole C. Robb, Jonathan M. Taylor, Amy Kent, Oliver J. Pambos, Barak Gilboa, Achillefs N. Kapanidis

## Abstract

Current virus detection methods often take significant time or can be limited in sensitivity and specificity. The increasing frequency and magnitude of viral outbreaks in recent decades has resulted in an urgent need for diagnostic methods that are facile, sensitive, rapid and inexpensive. Here, we describe and characterise a novel, calcium-mediated interaction of the surface of enveloped viruses with DNA, that can be used for the functionalisation of intact virus particles via chemical groups attached to the DNA. Using DNA modified with fluorophores, we have demonstrated the rapid and sensitive labelling and detection of influenza and other viruses using single-particle tracking and particle-size determination. With this method, we have detected clinical isolates of influenza in just one minute, significantly faster than existing rapid diagnostic tests. This powerful technique is easily extendable to a wide range of other enveloped pathogenic viruses and holds significant promise as a future diagnostic tool.

## INTRODUCTION

Infectious diseases caused by viruses represent a huge global public health concern, causing many thousands of deaths annually. The ability to efficiently and rapidly functionalise the surface of virus particles plays a major role in their study and characterisation. Virus functionalisation is not only important for viral manipulation, isolation and quantification, but importantly, can also allow viruses to be detected and identified. Early detection of viruses helps provide targeted therapy, lengthens the treatment window, reduces treatment costs and morbidity, helps prevent transmission, and leads to more efficient disease management and control.

Traditionally, the gold standard for viral detection has been virus culture in eukaryotic cell lines, which takes significant time (7-10 days) (reviewed in 1). Alternative nucleic acid amplification techniques such as RT-PCR are also commonly used; these assays are highly specific and sensitive, but can take several hours to obtain a result (reviewed in 1). Antigen-based rapid diagnostic tests are available for a small subset of viruses such as influenza and respiratory syncytial virus (RSV) and are relatively quick (<30 minutes), but limited by variable detection sensitivity (2). Diagnostic methods based on the direct quantification and identification of intact virus particles, such as flow cytometry (3, 4) have been used to quantify a variety of viruses with differing morphologies (reviewed in 5). These techniques make use of fluorescent dyes that bind to the viral genome (6) or proteins (7), all of which require a minimal 30 minute incubation period for efficient virus labelling prior to virus quantification. Negative stain transmission electron microscopy (TEM) images have been used since the 1940s (8) as an important tool to image individual virus particles and quantitatively determine virus concentrations, but due to the high costs and amount of space required for a TEM instrument, this is still only available in certain facilities. Therefore, despite significant progress in the development of viral diagnostics there is still an urgent need for detection methods that meet the criteria of being easy to use, sensitive, specific, rapid and cheap.

In this study, we have established a novel method to label and modify enveloped viruses using calcium chloride (CaCl_2_) and DNA. By adding fluorophores to the DNA, we were able to efficiently image and characterize viruses by their size in a single-particle tracking assay. This approach allowed us to detect viruses more rapidly than the currently available tests (with our results being available within one minute), and worked on all enveloped viruses tested, including viruses present in clinical samples. No amplification or purification steps were required as viruses could be detected directly in complex solutions such as cell culture media, virus transport media and allantoic fluid. We were also able to immobilise modified viruses, directly assess aggregation of fluorescent virus particles, and count labelled viruses using a fluidics-based optical approach. We therefore provide a novel tool for the rapid functionalisation, detection and quantification of enveloped viruses.

## RESULTS

### Calcium chloride labelling and single-particle tracking as a novel method for virus detection

Our study began with a serendipitous discovery while investigating methods of fluorescently labelling intact influenza virus particles. We observed that brief incubations of virus particles with short, fluorescently-labelled, non-specific DNAs in the presence of high concentrations of CaCl_2_ generated bright fluorescent particles that diffused slowly in solution, and resembled labelled viral particles (Movie S1). The fluorescent particles were extremely bright, pointing to the presence of multiple fluorescent DNA molecules per particle, and diffused much more slowly than free DNAs, which, due to their small size (only a few nm), diffused so rapidly that their motion blurred out and merely contributed to the overall background of the sample.

To establish whether the slowly diffusing particles were indeed labelled viral particles, we studied their mobility and number using single-particle tracking (reviewed in 9). Fluorescently-labelled particles were observed using a widefield microscope with variable angle epifluorescence microscopy (VAEM) to reduce the background signal (10) (Fig. 1A) and single-particle tracking software was used to track the bright and slow-moving particles (Fig. 1B). By localizing visible peaks and associating nearby localizations in subsequent frames with each other, the two-dimensional paths of the diffusing particles were reconstructed for the duration of their visibility in the focal plane (~700nm thick). The diffusion coefficients of the tracked particles were then plotted on a histogram and the mean diffusion coefficient (D_m_) was used to give an estimate of the average diameter of the diffusing particles (*d*_st_) using the Stokes-Einstein relation (Fig. 1C and equation 5). Thus, the diffusion coefficients and Stokes’ diameters obtained from single-particle tracking experiments could be used as an observable for particle detection.

**Figure 1.**
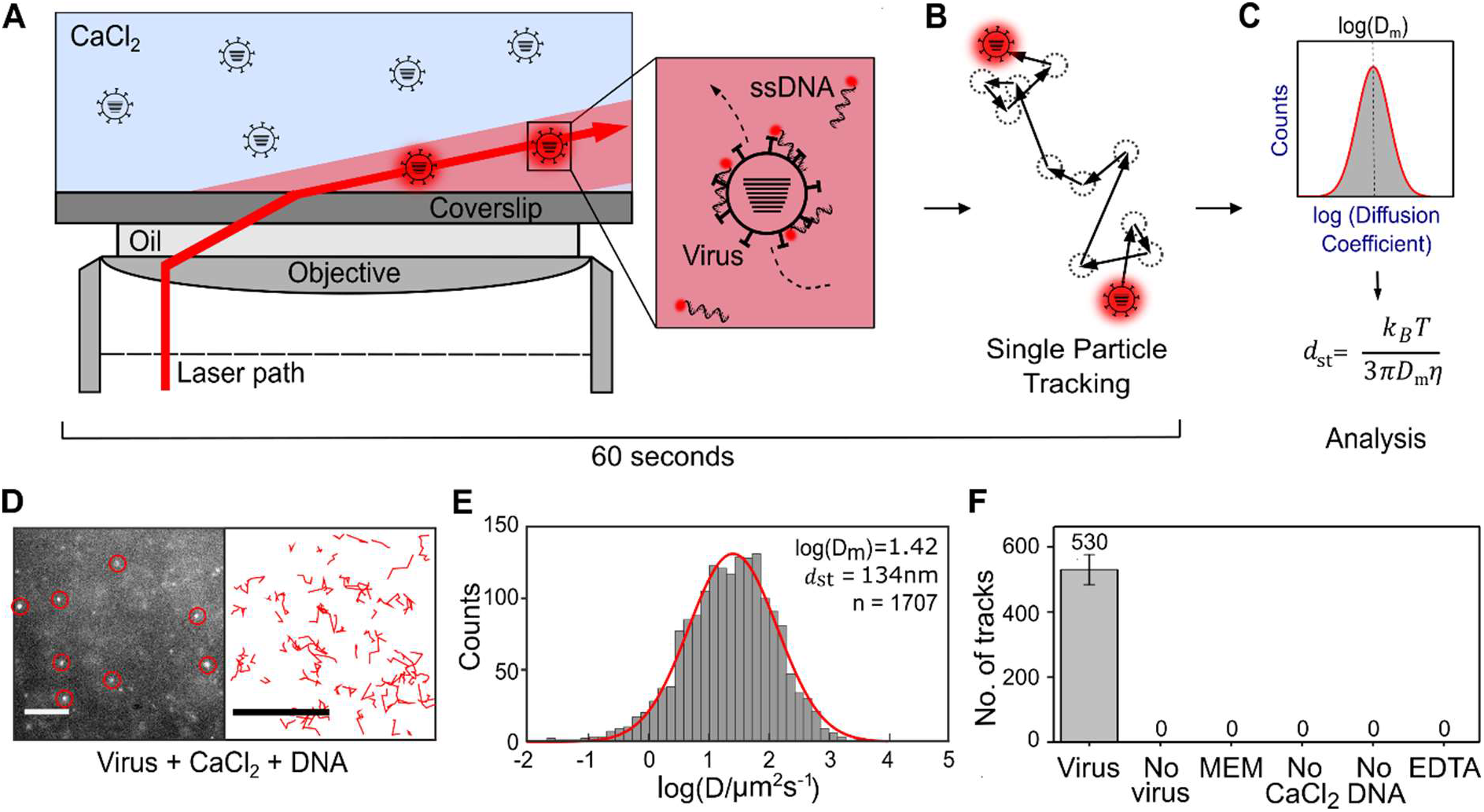
Detection of fluorescently-labelled influenza virus using single-particle tracking analysis. A) Schematic representation of the assay; where CaCl_2_ and fluorescent DNAs are used to non-specifically label diffusing virus particles. Multiply-labelled viruses are very bright, and move slowly, while free DNA is less bright and diffuses much more quickly. Samples are observed using wide-field variable-angle epifluorescence microscopy (VAEM). B) By localizing visible peaks and associating nearby localizations in subsequent frames with each other, the two-dimensional paths of labelled particles are reconstructed for the duration of their visibility in the focal plane. C) The diffusion coefficients of the tracked particles are plotted on a histogram and the mean diffusion coefficient (D_m_) is used to give an estimate of the average size of the diffusing molecules (*d*_st_) using the Stokes-Einstein relation. D) Representative field-of-view of fluorescently-labelled influenza virus diffusing in solution (left panel). A/Puerto Rico/8/1934 (H1N1) virus at a final concentration of 5.25×10^6^ PFU/mL was added to 0.65M CaCl_2_ and 1nM Atto647N-labelled DNA before being observed using a wide-field microscope. Scale bar 10μm. Red circles represent examples of localised particles. Representative tracks from the first 200 frames of the acquisition movie for fluorescently-labelled influenza virus diffusing in solution (right panel). Scale bar 10 μm. E) Diffusion coefficient histogram for A/Puerto Rico/8/1934. n = Number of tracks. F) Number of tracks observed in a 1000 frame acquisition in each condition. Negative controls where virus was replaced with water or minimal essential media (MEM), or CaCl_2_ or DNA were replaced with water, or after addition of 100mM of the calcium chelator EDTA to the imaging well. Error bars represent the standard deviation of 3 independent experiments.

Initially, we fluorescently labelled the H1N1 A/Puerto Rico/8/1934 (PR8) strain of influenza by adding 5.25×10^6^ PFU/mL PR8 to 0.65M CaCl_2_ and 1nM Atto647N-labelled single-stranded DNA (DNA 1, see methods), after which the sample was immediately transferred to the well of a glass slide placed on the microscope. Imaging of the sample showed the presence of multiple bright spots in the field-of-view (Fig. 1D, left panel). These spots were tracked (a 1000 frame acquisition at 30 Hz, taking 33 seconds) (Fig. 1D, right panel). Subsequent data analysis, taking less than 20 seconds, allowed diffusion coefficients to be calculated for each track (Fig. 1E). Fitting a Gaussian to the logarithm of the diffusion coefficients (D [μm^2^/s]), gave a mean diffusion coefficient log(D_m_) of 1.42, which provided a Stokes’ diameter of 134nm. The overall assay time took approximately 1 minute. Our results were also consistent with the tracking of commercial, fluorescently-labelled microspheres of a similar size (Fig. S1).

In order to verify the size and shape of a typical spherical influenza strain, we stained an A/Aichi/68 (X31) virus sample and imaged it using transmission electron microscopy (Fig. S2A&B). We measured the size (average of major and minor axes) (Fig. S2C) of 119 particles and obtained a broad distribution of virus sizes (Fig. S2D), with the largest number of particles (36) falling into the 120-140nm diameter range. This result was consistent with the results from our single-particle tracking assay, as well as with previous TEM or cryo-electron microscopy studies of influenza particles in the literature, which show that spherical strains generally consist of small spheres or ovoid particles of approximately 120nm in diameter (11–15).

To confirm that the signal in our assay was specific to fluorescently labelled PR8 virus particles, we replaced the virus with either water or minimal essential media (MEM) from mock-infected host cells. No signal was observed in the absence of virus (Fig. 1F & S3). Substitution of either the CaCl_2_ or the fluorescent DNA with water also resulted in no signal, indicating that they were both essential requirements for fluorescence labelling to occur. Interestingly, addition of the calcium chelator EDTA to the sample well during observation of fluorescently labelled virus particles immediately eliminated the signal, suggesting that the labelling is reversible (Fig. 1F & S3). Taken together, we have established that the bright, slow-moving particles we observe after incubation of influenza virus with CaCl_2_ and DNA are single, labelled viruses.

### Virus labelling requires calcium chloride, a fluorescent nucleic acid and a viral envelope

To investigate the nature of the ionic species required for fluorescence labelling to occur, CaCl_2_ was replaced with KCl, NaCl, MgCl_2_ or the cationic polyamine spermine, which has an overall ionic charge of +4. No tracks were detected when KCl or NaCl were used, while a very small number of tracks were detected when MgCl_2_ or spermine was used (an average of 6 and 25 tracks from a 1000-frame acquisition movie, respectively; Fig. 2A and S4A). In contrast, when CaCl_2_ was used, a large number of bright trackable particles were observed in the field of view (an average of 589 tracks from a 1000 frame acquisition movie; Fig. 2A and S4A). This suggests that labelling of virus particles using calcium chloride and DNA must involve a physical mechanism of interaction specific to CaCl_2_.

**Figure 2.**
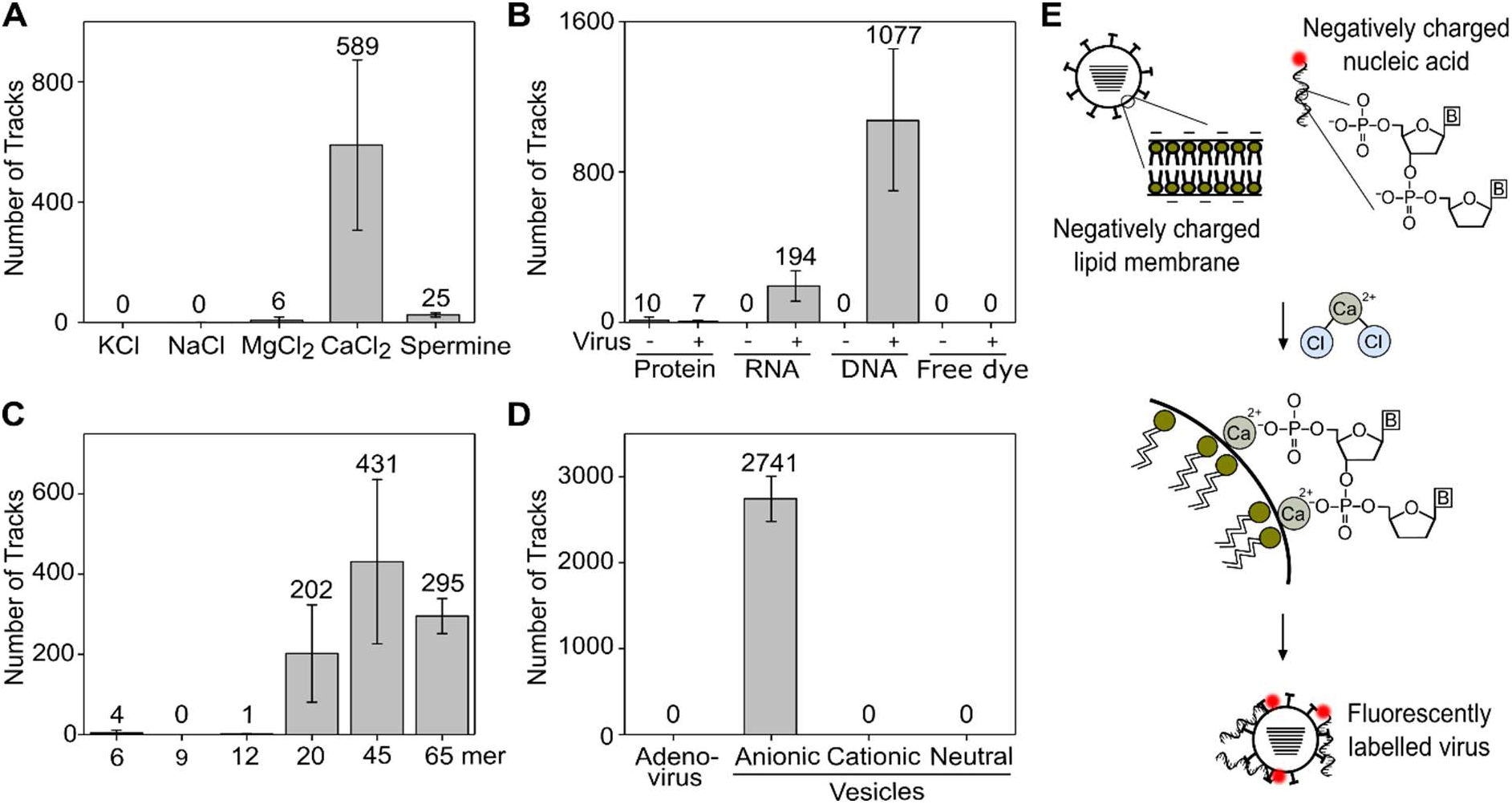
Virus labelling requires calcium chloride, a fluorescent nucleic acid and a viral envelope. A) Number of tracks observed in a 1000 frame acquisition when A/Puerto Rico/8/1934 (H1N1) virus at a final concentration of 26.25×10^6^ PFU/mL was added to 1nM Atto647N-labelled DNA and either 0.65M KCl, NaCl, MgCl_2_, CaCl_2_ or 0.32M spermine. The average number of tracks is indicated for each condition. Error bars represent the standard deviation of 3 independent experiments. B) Number of tracks observed in a 1000 frame acquisition when 1nM fluorescently-labelled DNA was replaced with 1nM Cy3B-labelled RNA, 1nM fluorescently-labelled protein (DNA polymerase) or 1nM free Atto647N dye. C) Number of tracks observed in a 1000 frame acquisition when A/Puerto Rico/8/1934 (H1N1) virus at a final concentration of 26.25×10^6^ PFU/mL was added to 1nM Atto647N-labelled DNA of different lengths and 0.65M CaCl_2_. D) Number of tracks observed in a 1000 frame acquisition when 3.3×10^11^ PFU/mL adenovirus was added to 1nM Atto647N-labelled DNA and 0.65M CaCl_2_ before being observed, or when virus was replaced with 200nm anionic, cationic or neutral charged lipid vesicles. E) Schematic model of calcium chloride-based fluorescent labelling of viruses. Enveloped viruses have an overall negative charge due to the phospholipids on the cell surface. DNA also has an overall negative charge due to multiple phosphate groups, B = base. Addition of CaCl_2_ provides divalent calcium cations that facilitate an interaction between the negatively charged polar heads of the viral lipid membrane and the negatively charged phosphates of the nucleic acid, possibly by binding two lipid molecules and a phosphate at the same time.

We also found that it was possible to label virus particles with either fluorescent single-stranded DNA or RNA, but not with a fluorescently-labelled protein (the Klenow fragment of DNA Polymerase; Fig. 2B and S4B), suggesting that efficient fluorescence labelling requires the presence of a nucleic acid. To eliminate the possibility that the nature of the dye was responsible for the labelling process, we attempted virus labelling and detection using unconjugated Atto647N dye not attached to a nucleic acid, which did not result in any fluorescent virus particles (Fig. 2B).

Next, in order to test the minimal length requirement of the nucleic acid required for efficient labelling, we tested fluorescently-labelled single-stranded DNAs of different lengths. We found that using short DNAs of 6, 9 or 12 nucleotides in length did not result in any appreciable virus labelling, while longer DNAs of 20, 45 or 64 nucleotides gave rise to bright fluorescent particles that could easily be tracked (Fig. 2C and S4C). Whilst use of the 45mer provided the highest number of tracks overall (Fig. 2C), the signal-to-noise improves with increasing length of DNA (Fig. S4C). Overall, our results suggest that fluorescent labelling via CaCl_2_ requires a nucleic acid greater than 12 nucleotides in length.

To study the requirement of a lipid membrane for efficient virus labelling, we tested a sample of nonenveloped adenovirus, which has a similar diameter to influenza, however no labelled particles were observed in the field-of-view and no tracks could be detected (Fig. 2D and S4D). Next, we calcium-labelled 200nm vesicles containing 25% anionic lipids (giving an overall negative charge), and found that we could observe and track bright particles diffusing in solution (Fig. 2D and S4D). When cationic or neutral 200nm vesicles were used in place of the anionic vesicles, no tracks were detected. Taken together, this strongly suggests that a negatively charged lipid bilayer is a necessary requirement for virus labelling to occur. We therefore propose a model for our labelling method that suggests that the Ca^2+^ ions derived from calcium chloride facilitate an interaction between the negatively charged polar heads of the viral lipid membrane and the negatively charged phosphates of the nucleic acid (see discussion) (Fig. 2E).

### Detection of clinical samples of influenza

In order to investigate the minimal virus concentration that we could reliably detect using our calcium-based labelling strategy, we tested increasing concentrations of virus. When virus was excluded from the sample (0 PFU/mL), no fluorescent particles were observed in the field-of-view; however, when PR8 virus was added, the number of fluorescent particles observed increased with increasing virus concentration (Fig. 3A). No tracks were detected in multiple acquisitions when virus was excluded from the sample, providing a value of zero for the ‘limit of blank’ (LOB); i.e., the highest number of tracks found when replicates of a blank sample containing no virus were tested (16). A small increase in the number of tracks was observed at low concentrations of virus (below 3.5×10^4^ PFU/mL), followed by an approximately linear increase for the higher concentrations tested (between 3.5×10^4^ and 26.2×10^4^ PFU/mL) (Fig. 3B). The ‘limit of detection’ (LOD; i.e., the lowest number of tracks likely to be reliably distinguished from the LOB and at which detection is feasible) was calculated as 1.86 tracks (16), corresponding to a minimal virus titre of 2.2×10^4^ PFU/mL (Fig. 3C).

**Figure 3.**
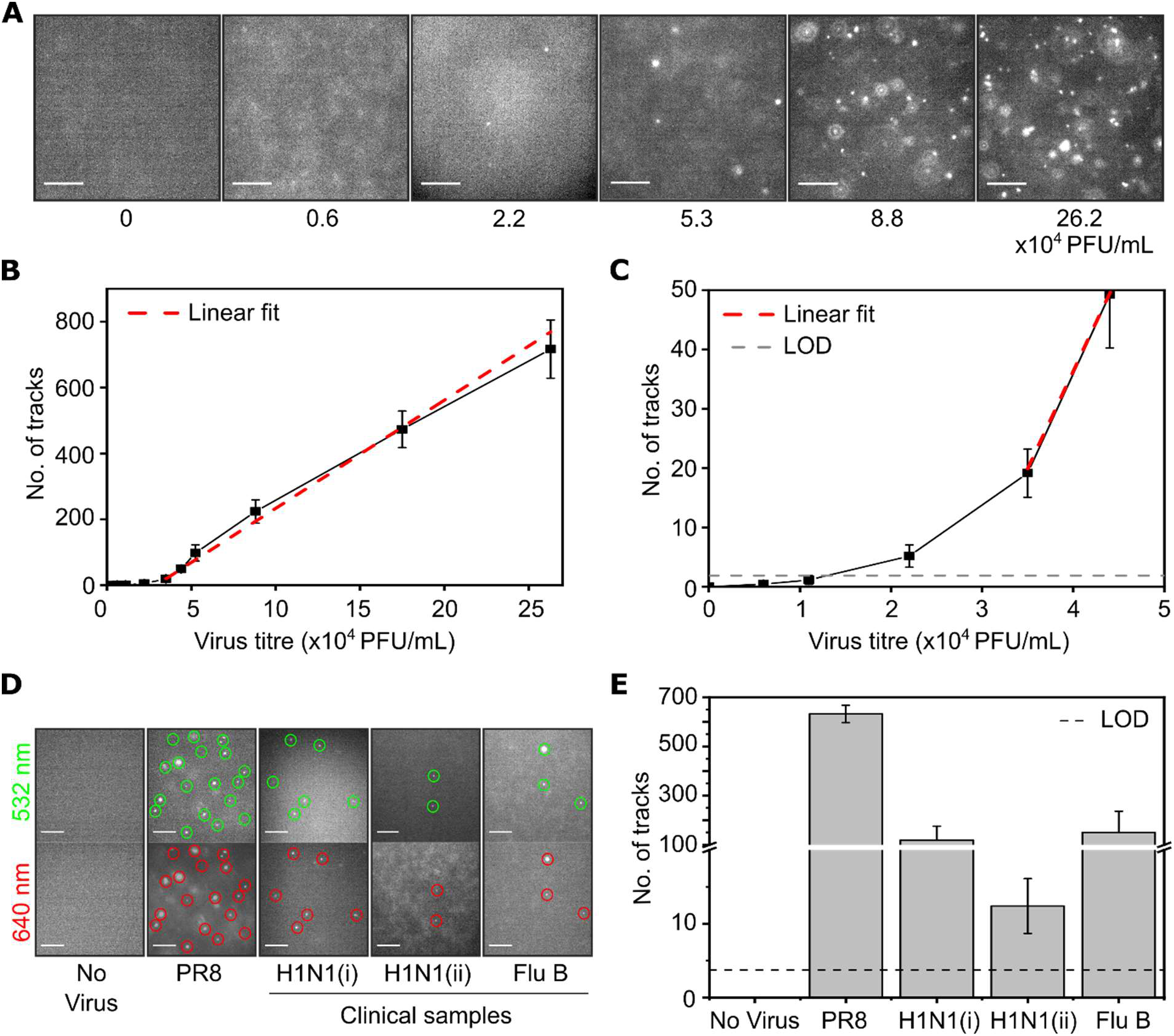
Detection of clinical samples of influenza. A) Representative fields-of-view of A/Puerto Rico/8/1934 (PR8) virus at a range of concentrations (described in figure) after being added to 0.65M CaCl_2_ and 1nM Atto647N-labelled DNA. Scale bar 10μm. B) Number of tracks observed in a 1000 frame acquisition at each concentration. Error bars represent the standard error of the mean from 7 independent experiments. C) Zoom in of number of tracks at lower concentrations. LOD = limit of detection. D) Representative fields-of-view of virus samples after being added to 0.65M CaCl_2_, 1nM DNA labelled with Atto647N, and 1nM DNA labelled with Cy3B. Virus strains used were PR8, two independent samples of A/California/07/2009 pandemic H1N1 virus (H1N1(i) and H1N1(ii)) and influenza B (Yamagata lineage) (Flu B). Scale bar 10μm. Green circles represent examples of localised particles in the 532 nm detection channel, red circles represent examples of localised particles in the 647 nm detection channel. B) Number of tracks (from the 647 nm channel) observed per 1000 frame acquisition for each virus sample. Error bars represent the standard deviation of 3 independent experiments.

Next, we examined clinical isolates of influenza. As we were expecting the virus concentration in clinical samples to be lower than laboratory-grown PR8, we detected viruses by adding two DNA species, labelled with either the red dye Atto647N (DNA 1) or the green dye Cy3B (DNA 2) simultaneously; this ensured that fluorescent viruses were concurrently labelled in both detection channels (that depend on excitation by 532- and 640-nm lasers; Fig. 4A and S5A&B), and thus were detected reliably. Using PR8 as an example, 86% of the detected virus particles were tracked in both channels simultaneously (Fig. S5C). As expected, when virus was excluded from the sample, no particles were observed in the field-of-view, and no tracks were detected (Fig 3D&E). Tracking of PR8 virus resulted in a high number of tracks (average of 632 from a 1000-frame acquisition; Fig. 3E). Two independent clinical samples of A/California/07/2009 pandemic H1N1 virus (H1N1(i) and H1N1(ii)) and influenza B (Yamagata lineage) (Flu B) were also tested, and resulted in an average 119, 12 and 150 tracks per 1000 frame acquisition (Fig. 3E). Although the number of detected viruses was lower in the clinical samples than in the laboratory-grown PR8 stock, this was still well above the LOD of our assay, clearly establishing that our detection assay could be useful in a clinical diagnostic setting.

**Figure 4:**
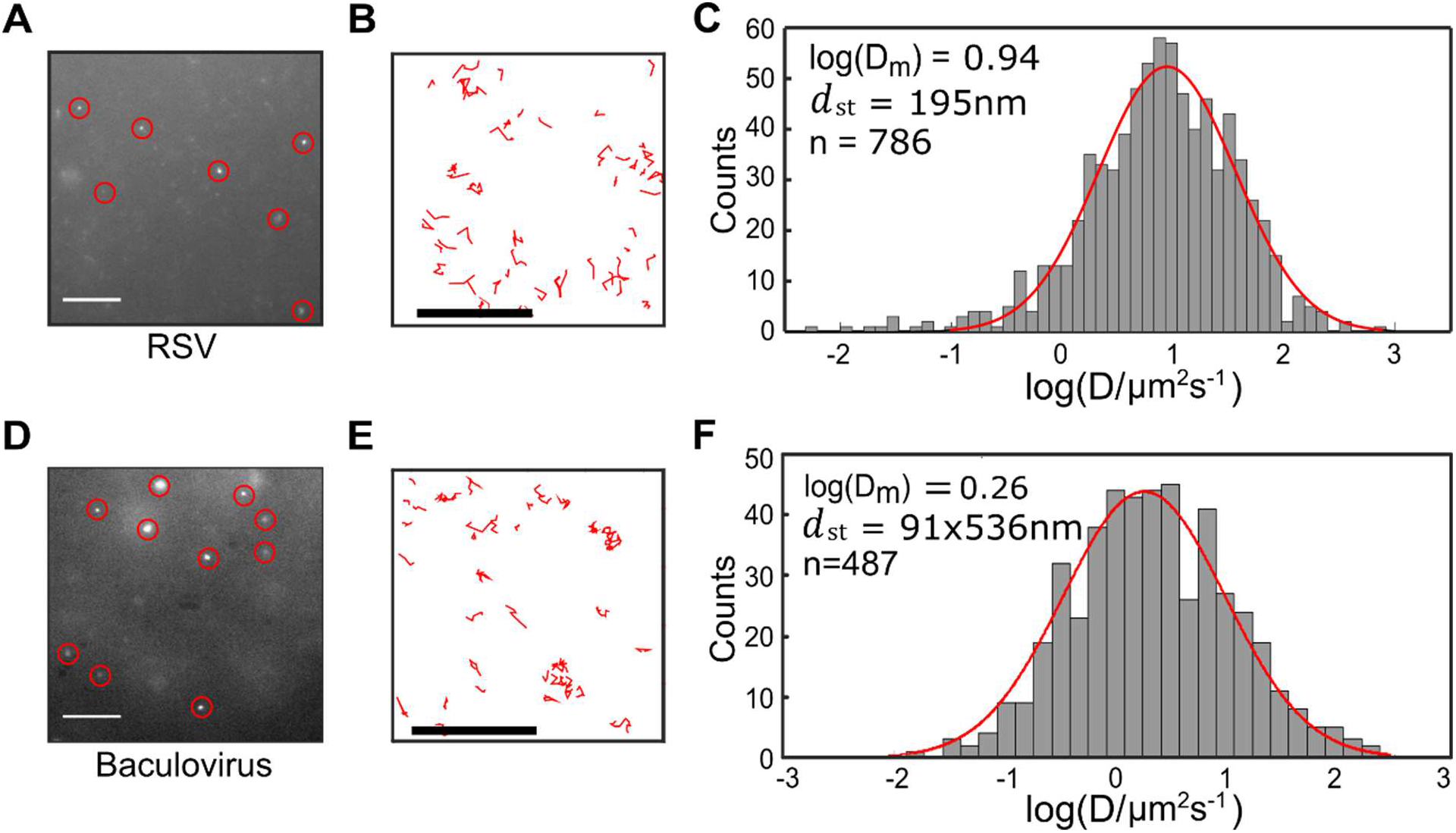
Detection of different viruses using fluorescent labelling and single particle tracking. A) Representative field-of-view of fluorescently-labelled respiratory syncytial virus (RSV) diffusing in solution. Virus at a final concentration of 7.0×10^2^ PFU/mL was added to 0.7M CaCl_2_, 1nM Atto647N-labelled DNA and 0.0025x trypsin before being observed using a wide-field microscope. Scale bar 10μm. Red circles represent examples of localised particles. B) Representative tracks from the first 200 frames of the acquisition movie. Scale bar 10 μm. C) Diffusion coefficient histogram for RSV. n = Number of tracks. D-F) As for A-C but for baculovirus. Baculovirus was added to 0.7M CaCl_2_ and 1nM Atto647N-labelled DNA before being observed using a wide-field microscope.

**Figure 5.**
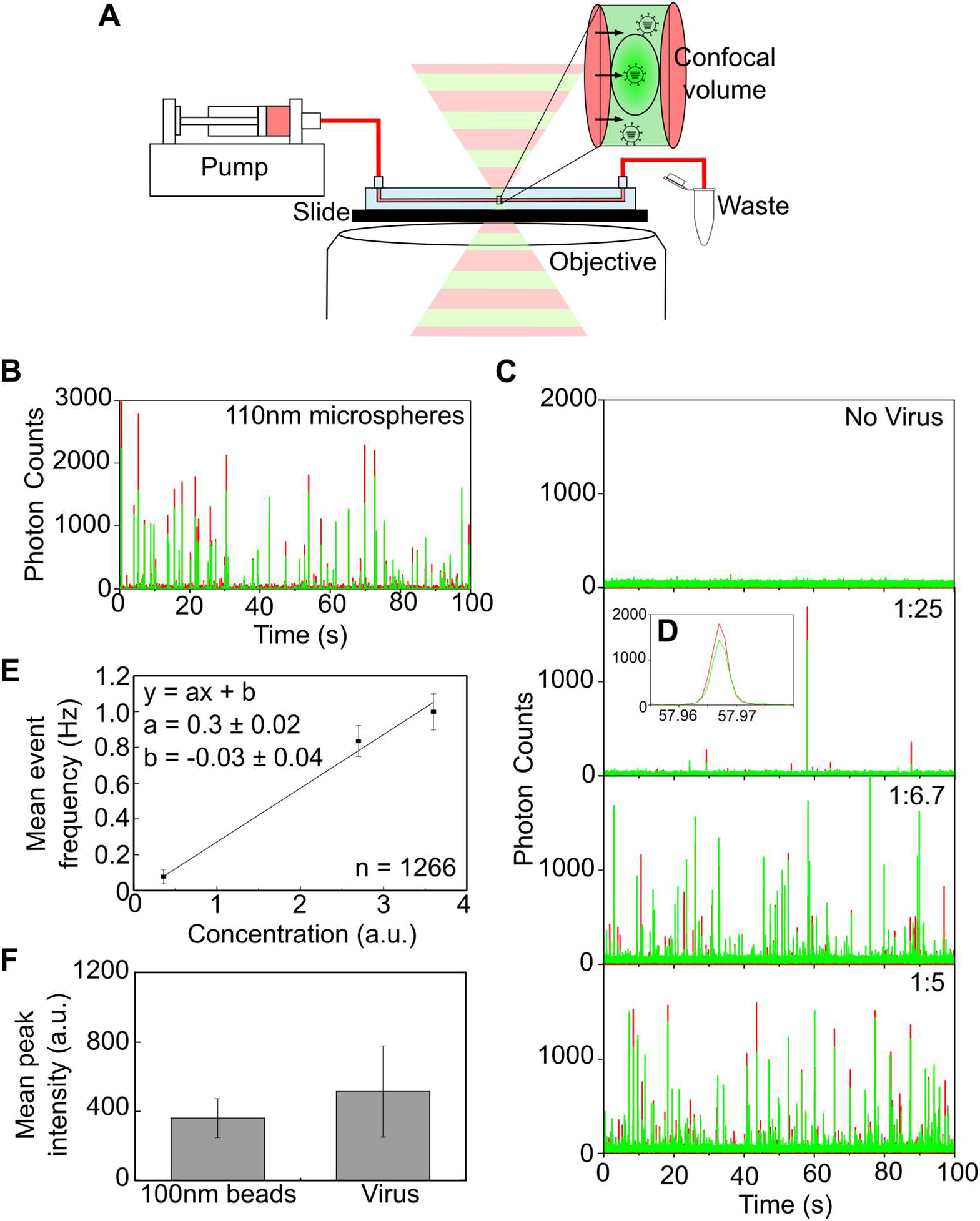
Non-specifically labelled influenza viruses are extremely bright, allowing rapid whole virus quantification. A) Schematic representation of the detection system used. Doubly-labelled viruses were flowed through a microslide channel (100μm high x 1mm wide) using a syringe infusion pump and illuminated using alternating red (647nm) and green (532nm) laser excitation at a modulation frequency of 10 kHz. Bursts of fluorescence corresponding to the passage of each virus particle through the confocal volume were split spectrally onto two avalanche photodiodes detecting red and green fluorescence. Custom-made LabVIEW software was used to register and evaluate the detector signal. B) Detection of 110nm fluorescent microspheres using the flow detection system. C) A/Puerto Rico/8/1934 virus at a final concentration of 5.25×10^6^ PFU/mL was doubly-labelled with 0.65M CaCl_2_ and 1nM Atto647N- and Cy3B-labelled DNA before being flowed through the microslide at increasing concentrations. D) Zoomed in image of a fluorescent burst corresponding to the passage of a virus particle through the confocal volume. E) Plot of mean number of fluorescent events occurring with increasing concentration of labelled virus. n = Number of bursts. F) Comparison of mean peak intensities of microspheres and virus particles. Error bars represent the standard deviation of the mean.

### Detection of multiple virus types and strains

In addition to the PR8 strain analysed previously (Fig. 1D-F), we were able to successfully detect influenza strains H1N1 A/WSN/1933 (WSN), H3N2 A/Aichi/68 (X31) and B/Florida/04/2006 (InfB) (Fig. S6). Viruses were detected in minimal essential media harvested from infected cells or allantoic fluid harvested from infected eggs, demonstrating the versatility of our assay for virus detection in different complex fluids. Interestingly, we found that different strains of influenza showed varying degrees of aggregation whilst diffusing in solution (for example, Fig. S6A,B,C shows several particles (circled) that visually appear brighter and larger than others in the field-of-view). Addition of trypsin reduced the aggregation of the virus particles, but did not eliminate it (Fig. S6). The larger Stokes’ diameters obtained for some virus strains suggested that we were observing a heterogeneous population of diffusing species consisting of some monomeric particles and some small aggregates. This theory was supported by simulations characterising the motions of populations of diffusing viruses with different sizes (Fig. S7). Using the X31 virus sample as an example, which had a log(D_m_) of 1.06 and mean size of 171nm, we found that a simulation movie consisting of 30% 100nm particles, 27% 200nm particles, 23% 300nm particles and 20% 400nm particles gave a mean size of 178nm when analysed (Fig. S7G). Comparison of the diffusion coefficient plots from the experimental and simulated data show an overall similarity in distribution and minimal differences in mean square deviation, while a comparison with simulated data from 100nm particles alone did not closely resemble the experimental data (Fig. S7G).

In addition to being applicable to the detection of a broad range of influenza strains, we tested our labelling method on two other enveloped virus types, the spherical human respiratory syncytial virus (RSV) and the Autographa californica nuclear polyhedrosis virus (AcNPV), a rod-shaped baculovirus of insects. Imaging and tracking of RSV gave a mean diffusion coefficient log(D_m_) of 0.94, which provided a mean diameter (*d*_st_) of 195nm (Fig. 4A-C and S8A-D), in good agreement with the size expected from the literature (17, 18). Baculovirus particles could also be efficiently detected (Fig. 4D&E and S8E-H). By modelling the viral particles as ellipsoids, with a characteristic rotational diffusion that was significantly smaller than the exposure time of our experiment (30ms), we assumed that the measured motion was isotropic (19). In this way, we were able to use the observed trajectories to calculate a diffusion coefficient histogram (equation 4, Materials and Methods) to give a mean diffusion coefficient log(D_m_) of 0.26 (Fig. 4F). By assuming a mean axial ratio of 5.9 (calculated by dividing a mean value of 325nm for the long axis of the baculovirus particle by 55nm for the short axis (19)), we obtained an approximate size of 91nm by 536nm for the baculovirus particles. Similarly to our experiments with influenza, over-estimation of virus size may result from aggregation. Nevertheless, our results have established the applicability of our virus detection methods to other enveloped viruses with a more complex geometry.

### Downstream applications of calcium-labelled virus detection

In order to demonstrate the applicability of our technique for virus counting (4, 5), we used CaCl_2_ and fluorescent DNA to label and directly quantify single virus particles in a simple fluidics system. Fluorescent viruses were pumped through a microslide channel and illuminated using a confocal microscope with alternating red and green laser excitation (Fig. 6A). As the fluorescent particles passed through the confocal volume, a burst of fluorescence was detected by two avalanche photodiodes detecting red and green fluorescence, and custom-written software was used to count the number of bursts above background (see materials and methods). As an initial proof-of-principle, we tested 110nm fluorescent microspheres, which showed frequent bursts of simultaneous red and green fluorescence as the spheres passed through the illuminated volume (Fig. 6B). Only low-level background fluorescence was detected when virus was excluded from the sample (Fig. 6C, top panel), however, when labelled PR8 virus was flowed through the channel, bursts of fluorescence showing simultaneous peaks of green and red fluorescence were observed (>90% of detected bursts showed both red and green labelling) (Fig. 6C second panel & 6D). The frequency of bursts increased linearly with increasing concentrations of virus (Fig. 6C, third and fourth panels and 6E).

When we quantified the mean intensity of the detected fluorescent bursts, we found that the intensities of the virus particles and the fluorescent microspheres were within error (standard deviation from multiple acquisitions) (Fig. 6F). By adding a biotin modification to the fluorescent DNA used for virus labelling we were able to surface-immobilise particles and obtain an estimate of the number of fluorescent DNAs per virus particle (Fig. S9A). By integrating the area of each intensity peak we obtained a mean intensity value of 3134 a.u. for the virus particles (Fig. S9B&C). By comparison, single immobilised fluorescent DNAs were significantly less bright than immobilised viruses, with a mean intensity value of 94 a.u. (Fig. S9D-F). This gives an average number of 33 DNA molecules per virus particle. Overall, the brightness of the virus particles arising from our labelling method suggests that it may be useful for viral detection methods based on direct quantification of intact virus particles, such as flow cytometry.

To demonstrate that our non-specific labelling approach could be combined with specific detection techniques, we used CaCl_2_ and DNA labelled with a green dye (Cy3B) to non-specifically detect immobilized virus particles, followed by staining of viruses with red fluorescent antibodies specific to the virus strain (Fig. S10A). Virus particles were visible in both the green (532 nm) and red (647 nm) channels, and the merged localisations showed many examples of co-localisation between the particles (Fig. S10B&C).

We also studied the ability of calcium-labelled influenza virus to bind to mammalian cells. After incubation of fluorescent PR8 virus with MDCK cells to allow adsorption, the cells were fixed, stained with DAPI to show cellular DNA in the cell nucleus, and imaged (Fig. S11). Small punctate spots were observed on the cell surface around the nuclei using 640nm excitation, suggesting that fluorescent virus particles were able to attach to host cells (Fig. S11A&B). These spots could not be seen when either PR8 (Fig. S11C), CaCl_2_ (Fig. S11D), or fluorescent DNA (Fig. S11E) were excluded from the experiment. Although it is unclear whether the labelled viruses enter the cells, their ability to bind to host cells suggests that labelled virus particles still maintain their structural integrity, and that the DNA does not completely obscure the viral haemagglutinin proteins required for host cell receptor binding on the virus surface.

## DISCUSSION

The interactions of calcium ions with lipid membranes have been probed by a variety of experimental methods, including fluorescent spectroscopy, dynamic light scattering and zeta potential measurements, in combination with molecular dynamics simulations (reviewed in 20). It is generally accepted that the presence of calcium rigidifies and orders lipid bilayers, by causing conformational changes in lipid headgroups, ordering of acyl chains, and lipid dehydration (reviewed in 20). Studies have demonstrated that calcium binds primarily to the phosphate groups of phospholipids, and simulations indicate that calcium is able to cluster phospholipid molecules via ion-bridges (21, 22), binding at least three lipid molecules concurrently (20, 22). It is plausible that in our experiments this stoichiometry is maintained, and one of the lipid molecules is replaced with a negatively charged phosphate group from the DNA/RNA (Fig. 2E). We cannot exclude the possibility that CaCl_2_ treatment also weakens the viral membrane and creates small pores or invaginations (23) that allow internalisation the nucleic acid, however the instantaneous loss of fluorescent signal upon EDTA addition suggests that the labelling is more likely to be external to the virus particle.

Our method for functionalisation of virus particles has several advantages over existing virus detection techniques. Our assay provides a simple and rapid method for quantitating whole virus particles, which is particularly important in the production of inactivated vaccines where virus titre cannot be estimated using traditional infectivity assays. The method also represents a quick way of assessing virus aggregation, and our results suggest that influenza viruses tend to aggregate independently in solution, in a strain-specific manner. We showed that viruses could be detected directly in complex biological fluids without the need for purification, that labelling was instantaneous and the signal from diffusing fluorescent viruses was readily detectable in just 1 minute. This timescale is more than an order of magnitude shorter than the conventionally used fluorescent dyes used to label viruses for flow cytometry. In addition, the technique is simple and cost-effective, and does not require expensive reagents. Only a small sample volume (1-20μl) is required, and virus can be detected without the need for an amplification step. We were able to detect influenza virus at a concentration of 2.2×10^4^ PFU/mL, and RSV virus at a concentration of 7.0×10^2^ PFU/mL. This detection limit can be lowered by taking longer acquisitions, with a small trade-off in overall assay time.

In this study, we have demonstrated the use of CaCl_2_ and DNA as a general biochemical tool for enveloped virus surface functionalisation, and demonstrated the feasibility of the approach in a rapid single-virus detection assay. The methods and analytical techniques developed here are applicable to the development of assays to study and detect many pathogenic viruses, and may be useful for the labelling or surface modification of any anionic lipid vesicles.

## METHODS

### Virus strains and clinical samples

H1N1 A/WSN/1933 (WSN), H1N1 A/Puerto Rico/8/1934 (PR8), H3N2 A/Udorn/72 (Udorn) and B/Florida/04/2006 (InfB) influenza viruses were grown in Madin-Darby bovine kidney (MDBK) or Madin-Darby canine kidney (MDCK) cells. The cell culture supernatant (Minimal Essential Media, Gibco) was collected and the viruses were titred by plaque assay. Titres of WSN, PR8, Udorn and InfB were 3.3×10^8^ plaque forming units (PFU)/mL, 1.05×10^8^ PFU/mL, 1.0×10^7^ PFU/mL and 2.1×10^8^ PFU/mL respectively. H3N2 A/Aichi/68 (X31) was grown in embryonated chicken eggs and titred by plaque assay (4.5×10^8^ PFU/mL). The A2 strain of RSV was grown in Hep-2 cells and titred by plaque assay (1.4×10^5^ PFU/mL). All influenza and RSV virus stocks were inactivated by shaking with 0.2% formaldehyde for 24 hours before use. Recombinant baculovirus derived from the *Autographa californica* nuclear polyhedrosis virus was produced using the Multibac system (24) and was of unknown titre. Chimpanzee adenovirus ChAdOx1-GFP was grown in HEK293 cells and titred by plaque assay (1.1×10^12^ PFU/mL) (25).

Clinical samples were collected as pharyngeal swabs (plastic shaft with Dacron tip) collected from patients presenting influenza-like illness. The swabs were inserted into tubes containing 2mL transport medium (GLY medium, Media Products, The Netherlands). The virus strain or subtype (A/California/7/2009 H1N1pdm09 and an influenza B clinical isolate of Yamagata lineage) was verified by Real Time RT-PCR.

### Fluorescent microspheres, DNA, RNA, protein and vesicles

Fluorescent microspheres with diameters of 110nm and 46nm were purchased from Life Technologies. The microspheres were diluted in water and sonicated for 15 minutes on ice prior to use.

Single-stranded oligonucleotides labelled with either Atto647N, Cy3B or Cy3 dyes were purchased from IBA (Germany). Unless specified otherwise the DNA sequence used was a 64mer labelled with Atto647N (DNA 1). For dual-colour labelling a second 64mer DNA labelled with Cy3B (DNA 2) was used. The RNA was a 34mer labelled with Cy3. Short DNAs used in Figure 2C were single stranded and labelled with either Atto647N or Cy3B. All sequences are provided in the supplementary information. The fluorescently-labelled protein was the Klenow fragment of E. coli DNA polymerase I (26), site-specifically labelled at position 550 with ATTO647N and 744 with Cy3B.

Lipid vesicles of 200nm in diameter were prepared as described previously (27), using extrusion with a 200nm pore size. Anionic lipid vesicles were composed of 75% 1,2-dioleoyl-sn-glycero-3-phosphocholine (DOPC) and 25% 1-palmitoyl-2-oleoyl-sn-glycero-3-phosphate (POPA), cationic vesicles were composed of 50% DOPC and 50% Ethyl-phosphocholine, and neutral vesicles were composed of soybean phosphocholine. Vesicles at a concentration of 10mg/mL were stored in 100mM KCl (pH 7.5), 50 mM MOPS and 1 mM MgCl_2_, before being diluted, labelled and imaged.

### Sample Preparation

Unless otherwise stated, virus stocks (typically 1-5μL) were diluted in 0.65M CaCl_2_ and 1nM fluorescently-labelled DNA in a final volume of 20μL (final virus concentrations are indicated in the figure legends). The sample was immediately placed in a well on a glass slide and imaged using variable angle epifluorescence microscopy (VAEM). The laser illumination was focused at a typical angle of 52° with respect to the normal, sufficiently above the surface of the glass slide to minimise background from unbound DNA settled on the surface (~3-15μm above surface). Typical acquisitions were 1000 frames, taken at a frequency of 30Hz, with laser intensities kept constant at 0.78 kW/cm^2^. In experiments where trypsin was added 0.5μL of a 0.05x stock of recombinant trypsin (TrypLE™ Express Enzyme, Thermo-Fisher Scientific) was added to the final sample volume of 20μL. In experiments where EDTA was added, a final concentration of 100mM EDTA was added directly to the well during imaging.

### Instrumentation

Single-particle tracking experiments were performed using wide-field imaging on a commercially available fluorescence Nanoimager microscope (Oxford Nanoimaging, https://www.oxfordni.com/). Briefly, a green (532nm) and a red (635nm) laser were combined using a dichroic mirror and coupled into a fibre optic cable. The fibre output was focused into the back focal plane of the objective (100x oil immersion, numerical aperture 1.4) and, for VAEM, displaced perpendicular to the optical axis resulting in a highly oblique subcritical incident angle on the sample to decrease background fluorophore excitation. Fluorescence emission was collected by the objective, separated into two emission channels and imaged onto a sCMOS camera (Orca flash V4, Hamamatsu).

### Data Analysis

For single-particle tracking analysis, the NanoImager software suite was first used to localize the fluorescent molecules in each frame by finding intensity peaks that were significantly above background, then fitting the detected spots with a Gaussian function. The NanoImager single-particle tracking feature was then used to map trajectories for the individual virus particles over multiple frames, using a pre-defined maximum step distance (the maximum distance that a particle may travel between frames) and nearest-neighbor exclusion radius. These parameters were used to maximise the track count; the maximum step distance was used to exclude particles that were diffusing much more quickly than the predicted size of virus particles (e.g. free dye) and the exclusion radius was used to exclude tracks which crossed over, which could lead to ambiguity in trajectory assignment. The 110nm and 46nm microspheres were used to calibrate the optimum parameters as a function of the expected distance travelled between frames and the spatial density of localisations. The maximum step distance was taken as

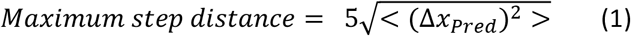

where < (Δ*x_Pred_*)^2^ > is the predicted mean squared displacement, and

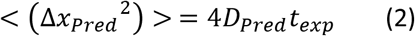

where *D_Pred_* is the predicted diffusion coefficient of a spherical particle of the expected size in the solution of interest and *t_exp_* is the exposure time (typically 30ms). The exclusion radius was taken as

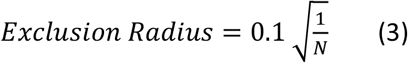

where *N* is the average number of particles per unit area, calculated by averaging the number of particles per unit area in each frame of a movie. The channel with the largest number of localisations was used to calculate the exclusion radius. In cases where the exclusion radius was smaller than the maximum distance, in samples with a high density of fluorescent particles for example, both parameters were set to the maximum distance. In the case of multicolour tracking, a particle observed in both channels was considered to be co-localized if simultaneous localisation in the red and green channels was separated by less than 200nm.

The trajectories were then used to calculate a diffusion coefficient for each track using the equation:

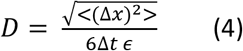

where *D* is the diffusion coefficient, <(Δ*x*)^2^> is the mean square displacement, Δ*t* is the change in time and ϵ is a correction factor to account for a non-zero exposure time (28). A histogram of the natural log diffusion coefficient for each track was then plotted using a custom-written MATLAB script. The histogram was fitted with a Gaussian function and the diffusion coefficient which corresponded to the curve peak was taken to be the sample mean (D_m_). The Stokes’ diameter (*d*_st_) was then calculated using:

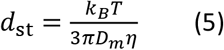

where T is the temperature (in Kelvin), D_m_ is the mean diffusion coefficient and *η* is the viscosity of the solution. The sample temperature was obtained from the microscope sensors and the viscosity of the solution was corrected for the presence of calcium chloride (29).

### Simulations

Movies simulating the unconfined diffusion of viruses were produced using existing software (30), with a number of modifications. For each virus particle, an initial position and time at which the particle entered the field-of-view was generated randomly, with the bounds of diffusion set to beyond the field-of-view to simulate unconfined diffusion. The position of each particle was then calculated at 100 sub-steps per frame to simulate the effects of motion blur, using track lengths that followed an exponential distribution. Point-spread-functions were calculated for each localisation and added to a matrix that formed the final movie, before random noise was added to simulate fluctuations in the background photon intensity. The simulated movies were then analysed using custom-written MATLAB software.

### Electron microscopy

H3N2 A/Aichi/68 (X31) virus was visualized by transmission electron microscopy at the Bioimaging Facility of the Sir William Dunn School of Pathology, University of Oxford. Virions were prepared as described previously (31). Briefly, they were fixed with 4% paraformaldehyde, adsorbed onto grids, negatively stained with 2% aqueous uranyl acetate and imaged in a Tecnai 12 transmission electron microscope (FEI, Eindhoven) operated at 120 kV.

### Virus Counting

A custom-built confocal microscope was used for single-particle virus counting experiments (32–34). Fluorescent microspheres with a diameter of 110nm or fluorescently-labelled A/Puerto Rico/8/1934 virus was flowed through a microslide channel (100μm high x 1mm wide; Ibidi GmbH) using a syringe infusion pump (Harvard Apparatus, Pump 11 Elite) and illuminated using alternating red (647nm) and green (532nm) laser excitation at a modulation frequency of 10 kHz. Bursts of fluorescence corresponding to the passage of each virus particle through the confocal volume were split spectrally onto two avalanche photodiodes detecting red and green fluorescence. Fluorescence data recorded by the avalanche photodiodes was acquired using a custom-written LabVIEW virtual instrument into the single molecule (.sm) format, and converted to plain text using a second virtual instrument. Analysis of the raw time series data was performed using custom-written MATLAB code, which implemented a threshold-based sliding window algorithm to detect events, combined with a Gaussian peak fitting process to identify the height, width, and location of individual peaks. Using a trigger level, re-arming level, minimum event duration, maximum event intensity, and baseline input by the user, the procedure first subtracted the baseline. The trace was then scanned point-by-point, building a map of time points for which the fluorescence intensity exceeds the trigger level without re-crossing the re-arming threshold. To minimise errors in the Gaussian peak fitting process for short-lived events, data were automatically extracted from either side of these events to ensure that a minimum of 10 data points were available to the peak fitting algorithm. Data from each event were fitted using a Gaussian peak fitting procedure, outputting the peak intensity, duration, and position within the trace of each fluorescent event. In rare instances, where the expansion of the event window caused overlap with neighbouring events, fitting errors may occur. To address this issue, the procedure identified overlapping regions, which were then merged. The peak fitting process then identified only the largest peak, and discarded the smaller peaks. Error checking was then performed, removing events that exceeded the maximum expected event duration, and intensity. Descriptive statistics for detected peaks were output for further analysis and presentation in Origin.

### Intensity profile Analysis

A/Puerto Rico/8/1934 (H1N1) virus particles were fluorescently labelled by incubation with 1nM biotin-conjugated DNA labelled with Atto647N, before being immobilised on the surface of a pegylated glass slide treated with neutravadin. The slide surface was imaged using TIRF (total internal reflection fluorescence) with the laser illumination focused at a typical angle of 53° with respect to the normal. Intensity profiles from 20 pixel slices of the fields of view were taken using the built-in ‘plot profile’ feature on ImageJ. The intensity peaks were fitted using the ‘fitpeaks’ function in MATLAB, the local background was subtracted, and the peaks were integrated to give an estimate of the intensity of the peak. Multiple particles were analysed and the mean or median peak area of the resulting histogram was taken as the average particle intensity, which was compared between virus particles and free DNA.

### Antibody and cell staining

Virus was incubated with 0.65M CaCl_2_ and 1nM biotinylated Cy3B-labelled DNA before being immobilized via neutravidin on the surface of a pegylated glass slide. Viruses were fixed in 4% paraformaldehyde and permeabilised with 0.5% Triton-X-100, before being incubated with primary and secondary antibodies. A/Udorn/72 virus was stained with an anti-NA primary antibody and Alexa647-labelled secondary antibody and A/WSN/33 virus was stained with an anti-NP primary antibody and Alexa647-labelled secondary antibody. Localisations in each channel were identified using the NanoImager software suite.

MDCK cells were grown on 13mm coverslips in 24-well plates before being infected with A/Puerto Rico/8/1934 virus at a multiplicity of infection of ~20×10^6^ PFU/mL. The virus was incubated with 0.65M CaCl_2_ and 1nM Atto647N-labelled DNA before being added to the cells. The cells were incubated for 1 hour at 37°C to allow viruses to adhere before being fixed for 15 minutes in 4% paraformaldehyde and mounted onto glass microscope slides with Mowiol containing DAPI. Cells were viewed using an Olympus Fluoview FV1200 microscope and images processed with ImageJ.

### Statistical analysis

The Limit of Blank (LOB) and Limit of Detection (LOD) were calculated as described in (16). The LOB was defined as the highest concentration found when replicates of a blank sample containing no virus were tested, and was calculated using:

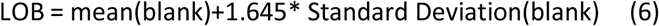

The LOD was defined as the lowest virus concentration reliably distinguished from the LOB and at which detection was feasible, determined by utilizing both the measured LOB and test replicates of a sample known to contain a low concentration of virus (0.6×10^4^ PFU/mL).

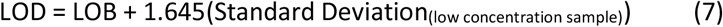

## ACKNOWLEDGEMENTS

We thank Professor Ervin Fodor and Dr. David Bauer at the Sir William Dunn School of Pathology, Oxford for the WSN, PR8 and InfB influenza virus stocks, Michael Bennett at The Francis Crick Institute for the X31 influenza virus stock, the Oxford Vaccine Group for the RSV stocks, Dr. Alison Turner and Professor Sarah Gilbert at The Jenner Institute for the adenovirus stock, Dr Maria Evangelidou and Dr Alexios-Fotios Mentis at the National Influenza Reference Center of Southern Greece, Hellenic Pasteur Institute, for the clinical influenza samples, Errin Johnson at the Bioimaging Facility of the Sir William Dunn School of Pathology, University of Oxford for the transmission electron microscopy, Dr. Robert Ishmukhametov at the University of Oxford for providing the lipid vesicles, Rebecca Andrews for providing the short DNA oligos, Thomas Jollans and Fraser Jenkings for their contribution in the early development of this work and the Micron Imaging Facility, University of Oxford, for access to microscopes. This work was supported by Wellcome Trust grant 110164/Z/15/Z and Medical Research Council (MRC) grant MR/N010744/1 (to A.N.K.), and a Royal Society Dorothy Hodgkin Research Fellowship DKR00620 (to N.C.R.).

## AUTHOR CONTRIBUTIONS

N.C.R., J.M.T, A.K. and A.N.K. designed and carried out experiments, analysed data and interpreted results. O.J.P wrote analysis software and analysed results. B.G. wrote analysis software. N.C.R., J.M.T. and A.N.K. wrote the manuscript. All authors reviewed and approved the final manuscript.

