## Supplemental files for "Rapid functionalisation and detection of viruses via a novel Ca^2+^-mediated virus-DNA interaction"

**Supplementary Material**

**Movie S1. Movie of fluorescently labelled virus particles diffusing in solution.** A/Puerto Rico/8/1934 (H1N1) virus at a final concentration of $5.25 \times 10^6$ PFU/mL was added to 0.65M $CaCl_2$ and 1nM Atto647N-labelled DNA before being observed using a wide-field microscope. The movie shows a 1000 frame acquisition, taken at a frequency of 30Hz, with laser intensities kept constant at 0.78 kW/cm$^2$. Scale bar is 10μm.

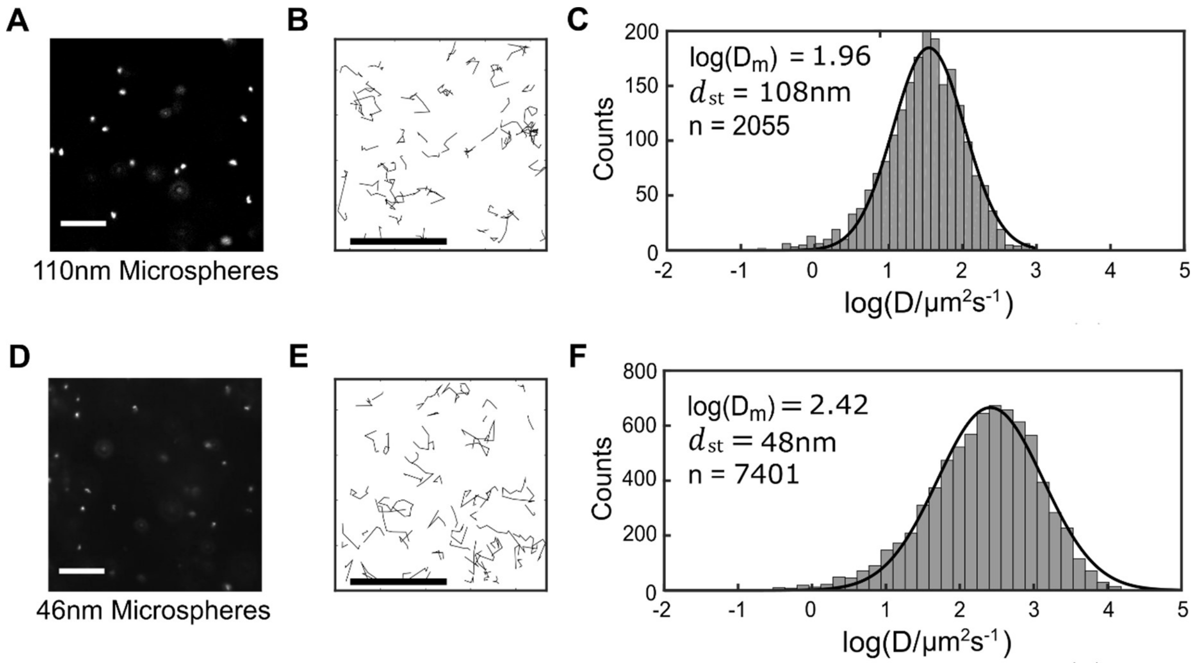


**Supplementary Figure 1. Differentiation between fluorescent microspheres of different sizes using single-particle tracking analysis.** A) Representative field-of-view of 110nm fluorescent microspheres diffusing in water. Scale bar 10 μm. B) Representative tracks from the first 200 frames of the acquisition. Scale bar 10 μm. C) Diffusion coefficient histogram for the 110nm microspheres. n = Number of tracks. D) Representative field-of-view of 46nm fluorescent microspheres diffusing in water. Scale bar 10 μm. E) Representative tracks from the first 200 frames of the acquisition. Scale bar 10 μm. F) Diffusion coefficient histogram for the 46nm microspheres. n = Number of tracks.

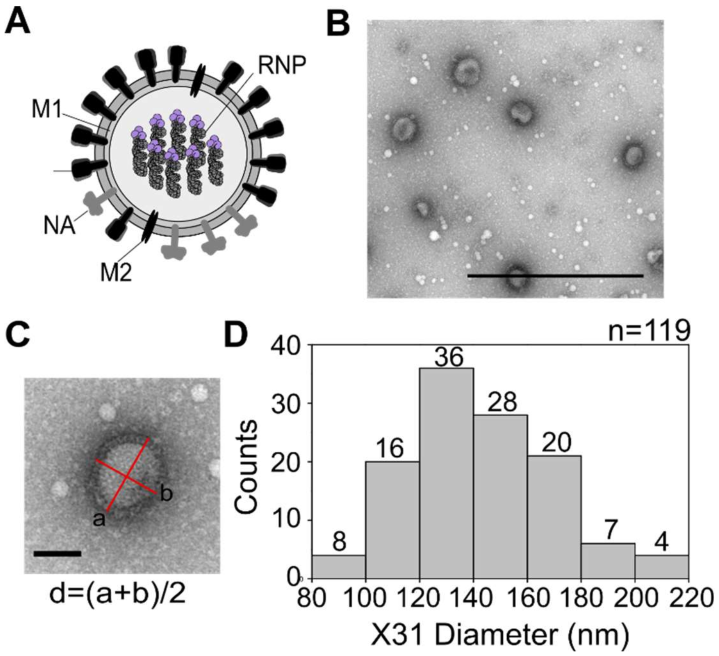


**Supplementary Figure 2. Electron microscopy measurement of influenza virus particle size.** A) Schematic representation of an influenza particle. Virus particles have three transmembrane proteins (HA, NA and M2), beneath which lies a layer of matrix protein (M1), which encloses the eight genomic ribonucleoproteins (RNPs). B) Representative electron micrograph image of negatively stained X31 influenza particles. Scale bar 1 µm. C) The mean diameter of each virus particle was calculated by taking the average between major and minor axes. Scale bar 0.1 µm. D) Histogram of the mean diameter of the X31 virus. Number of virus particles counted (n) = 119.

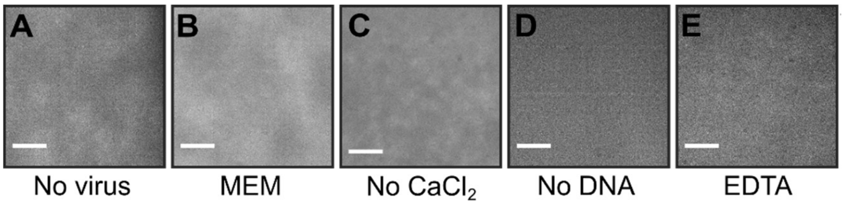


**Supplementary Figure 3 (related to Figure 1). Fluorescent particles are only detected in the presence of virus, $CaCl_2$ and DNA.** Negative controls where virus was replaced with A) water or B) minimal essential media. Negative controls where C) $CaCl_2$ or D) DNA were replaced with water. E) Addition of 100mM of the calcium chelator EDTA to the imaging well resulted in the loss of signal**.** Scale bar 10μm

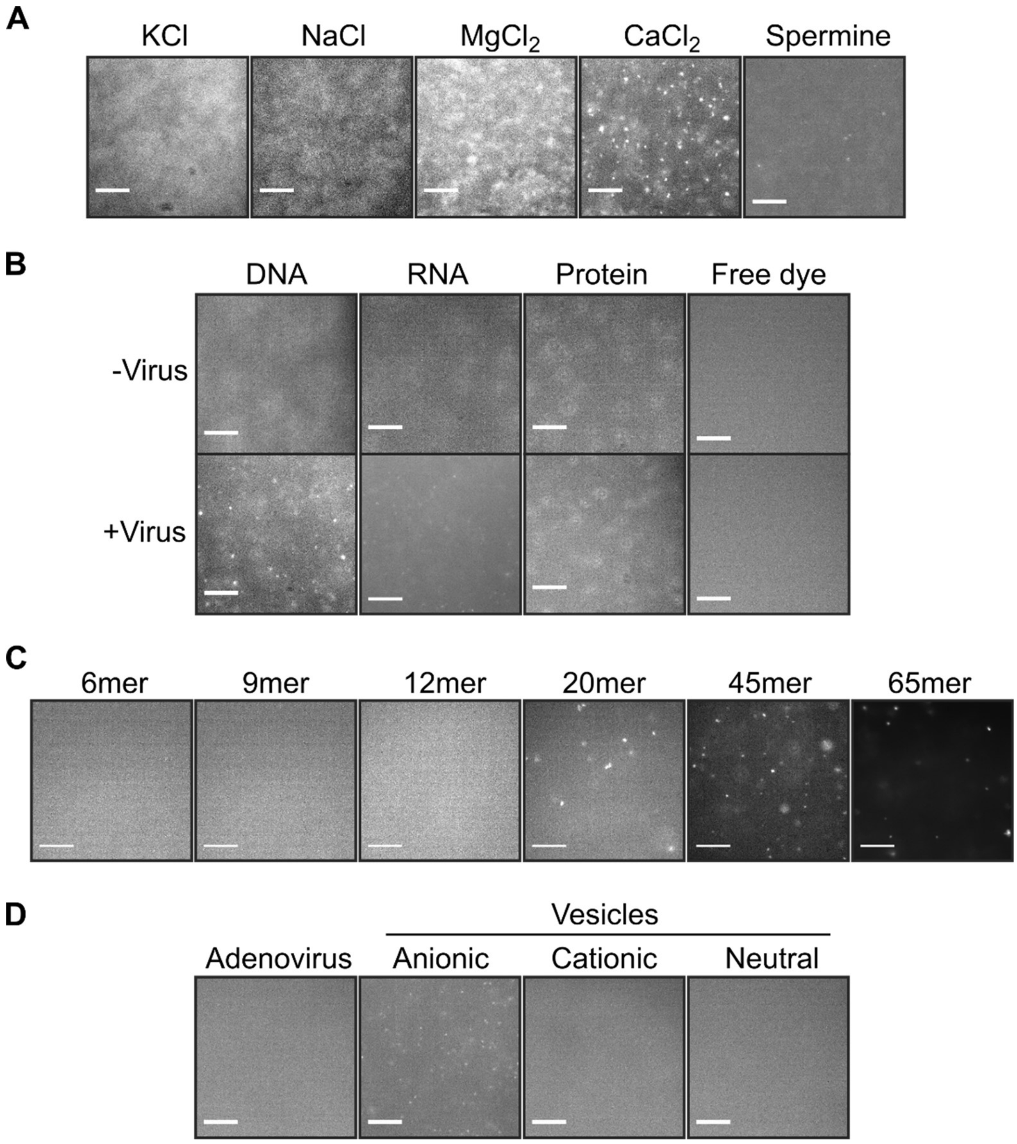


**Supplementary Figure 4 (related to Figure 2). Virus labelling requires calcium chloride, a fluorescent nucleic acid and a viral envelope.** A) Representative fields-of-view when A/Puerto Rico/8/1934 (H1N1) virus at a final concentration of $26.25 \times 10^6$ PFU/mL was added to 1nM Atto647N-labelled DNA and either 0.65M KCl, NaCl, $MgCl_2$, $CaCl_2$ or 0.32M spermine. B) Representative fields-of-view when 1nM fluorescently-labelled DNA was replaced with 1nM fluorescently-labelled RNA, 1nM fluorescently-labelled protein (DNA polymerase) or 1nM free Atto647N dye. C) Representative fields-of-view when A/Puerto Rico/8/1934 (H1N1) virus at a final concentration of $26.25 \times 10^6$ PFU/mL was added to 1nM Atto647N-labelled DNA of different lengths and 0.65M $CaCl_2$. Scale bars are 10 µm. D) Representative fields-of-view when $3.3 \times 10^{11}$ PFU/mL adenovirus was added to 1nM Atto647N-labelled DNA and 0.65M $CaCl_2$ before being observed, or when virus was replaced with 200nm anionic, cationic or neutral charged lipid vesicles. Scale bars are 10 µm.

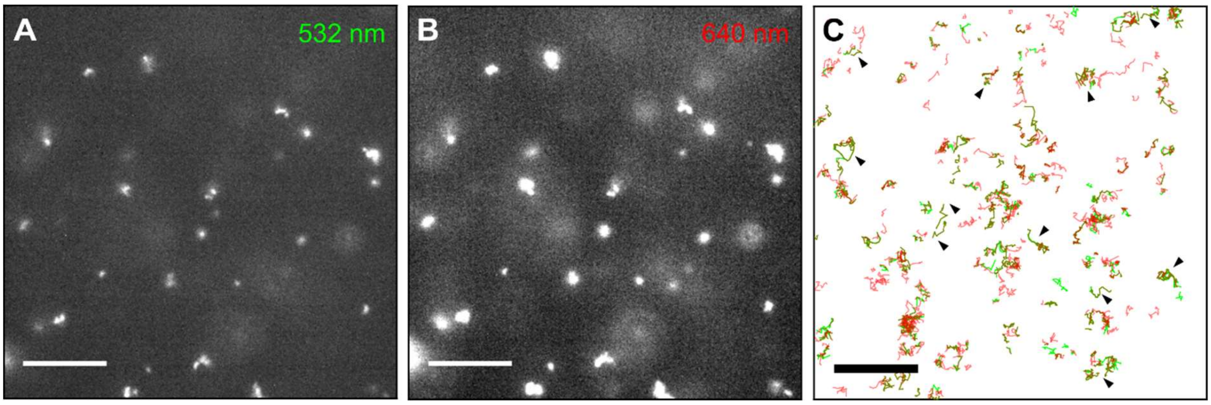


**Supplementary Figure 5 (related to Figure 3). Dual-colour labelling and virus tracking with different dyes.** A&B) Representative fields-of-view (from frame 1) of doubly-labelled influenza viruses diffusing in solution, with A) 532 nm laser illumination and B) 640 nm laser illumination. A/Puerto Rico/8/1934 (H1N1) virus at a final concentration of $5.25 \times 10^6$ PFU/mL was added to 0.65M $CaCl_2$, 1nM DNA labelled with Atto647N, and 1nM DNA labelled with Cy3B, before being observed using a wide-field microscope. Scale bar 10µm. C) Representative tracks from doubly-labelled viruses. Black arrows show examples of trajectories from double-labelled viruses. Scale bar 10 µm.

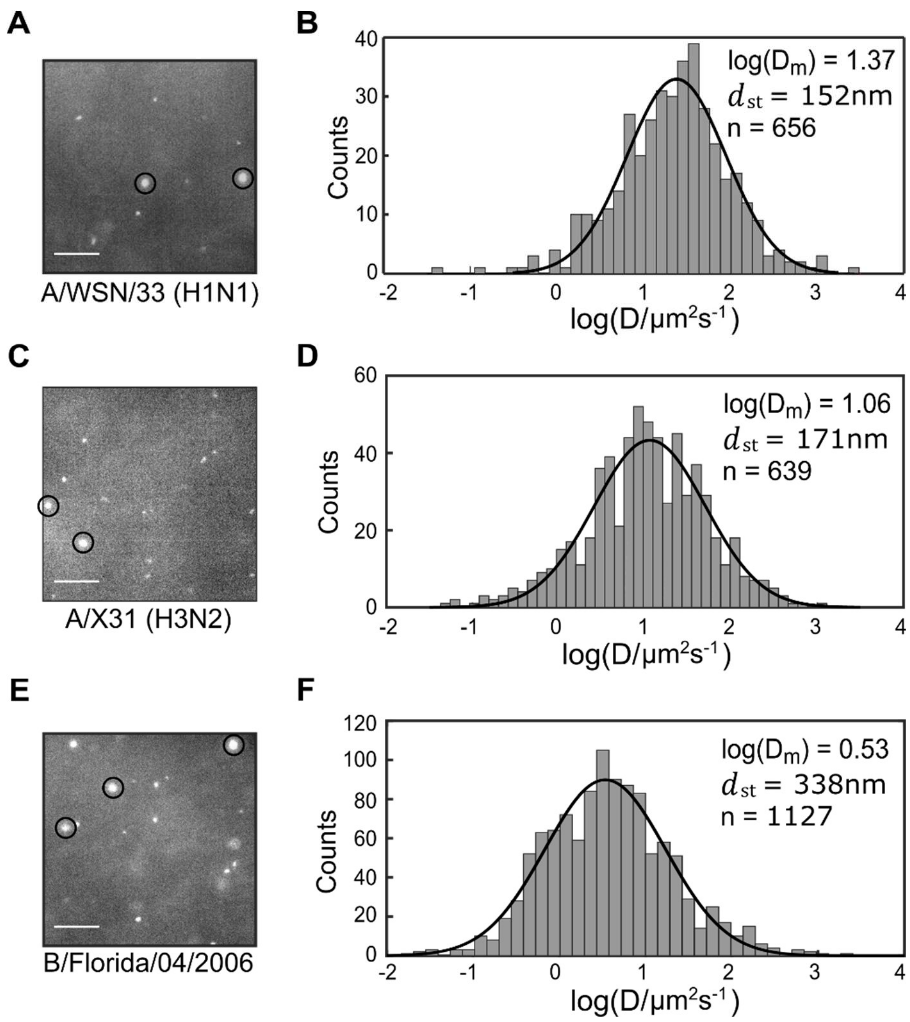


**Supplementary Figure 6. Detection of different influenza strains using fluorescent labelling and single particle tracking.** A) Representative field-of-view of fluorescently-labelled A/WSN/33 influenza virus diffusing in solution. Virus at a final concentration of 49.5x$10^6$ PFU/mL was added to 0.65M $CaCl_2$, 1nM Atto647N-labelled DNA and 0.0025x trypsin before being observed using a wide-field microscope. Scale bar 10µm. B) Diffusion coefficient histogram for A/WSN/33. n = Number of tracks. C&D) As for A&B but for A/X31 influenza virus diffusing in solution. Virus at a final concentration of 22.5x$10^6$ PFU/mL was added to 0.65M $CaCl_2$, 1nM Atto647N-labelled DNA and 0.0025x trypsin before being observed using a wide-field microscope. Scale bar 10µm. E&F) As for A&B but for B/Florida/04/2006. Virus at a final concentration of 3.15x$10^6$ PFU/mL was added to 0.65M $CaCl_2$, 1nM Atto647N-labelled DNA and 0.0025x trypsin before being observed using a wide-field microscope. Circles represent larger aggregates of virus particles.

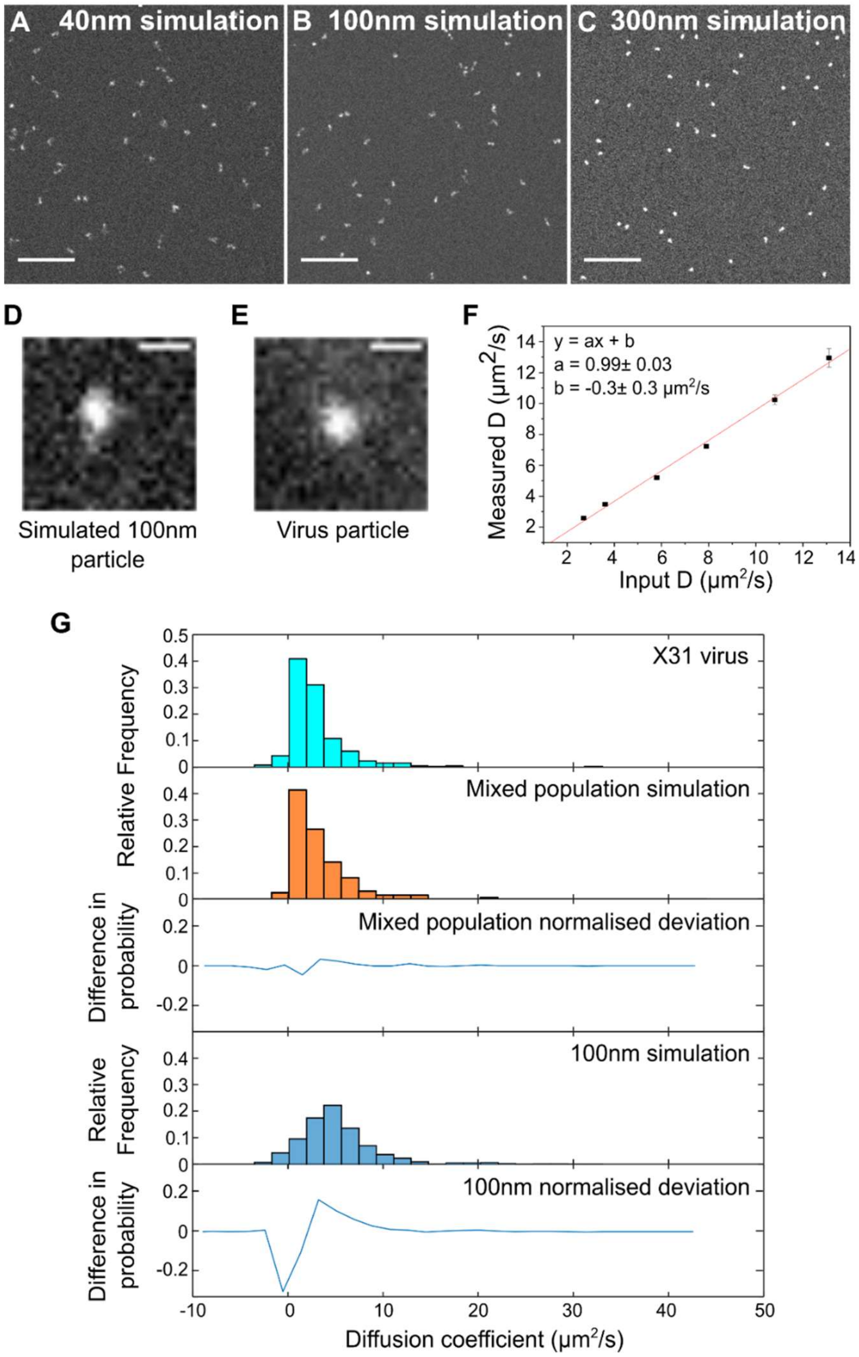


**Supplementary Figure 7 (related to Figure S6). Simulations to investigate virus aggregation by characterising the observed motions of a diverse population of diffusing particles of different sizes.** A) Representative field-of-view from a simulation movie of 40nm particles diffusing within a 400x400 pixel area. Scale bar 10 µm. B) Field-of-view from a simulation movie of 100nm particles. Scale bar 10 µm. C) Field-of-view from a simulation movie of 300nm particles. Scale bar 10 µm. D) Representative 100nm particle from a simulated movie. Scale bar 5 µm. E) Zoomed in image of a fluorescently-labelled A/WSN/33 influenza virus diffusing in solution. Virus at a final concentration of $49.5 \times 10^{6}$ PFU/mL was added to 0.65M $CaCl_2$, 1nM Atto647N-labelled DNA and 0.0025x trypsin before being observed using a wide-field microscope. Scale bar 5 µm. F) Simulation input diffusion coefficients and the values measured by the software were consistent within error. G) Diffusion coefficient plots from the experimental (X31 virus) and simulated data (mixed population of 100nm, 200nm, 300nm and 400nm particles, or 100nm particles only) and differences in mean deviation.

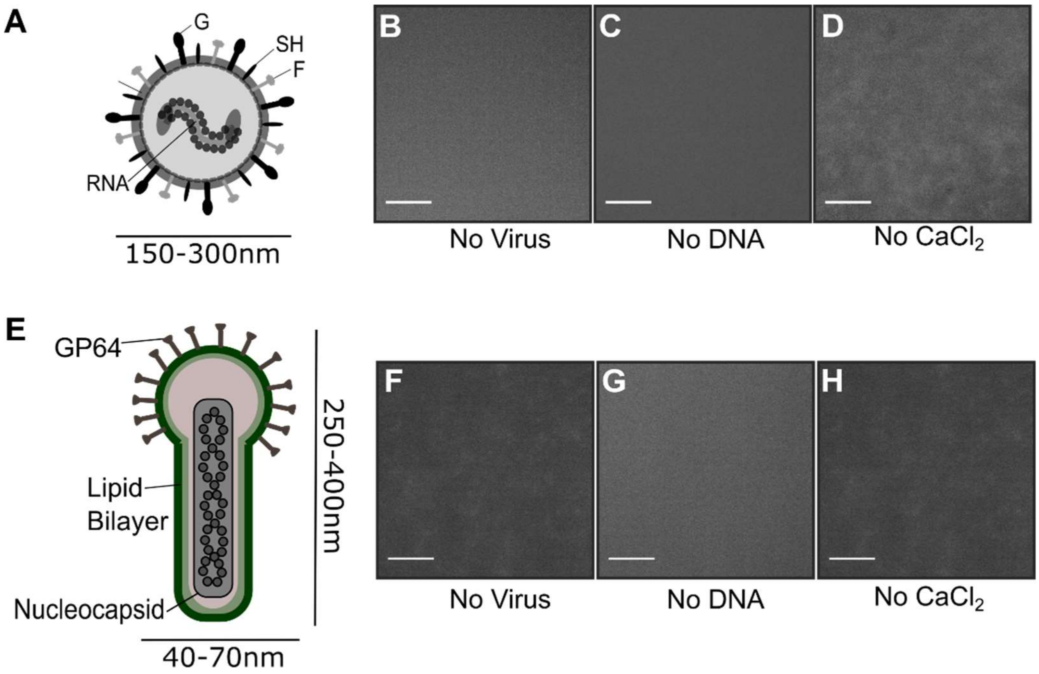


**Supplementary Figure 8 (related to Figure 4). Calcium chloride can be used to fluorescently label respiratory syncytial virus (RSV) and baculovirus.** A) Schematic representation of an RSV particle. Virus particles have three transmembrane proteins (attachment (G) protein, fusion (F) protein and small hydrophobic (SH) protein), beneath which lies a layer of matrix protein (M), which encloses the single-stranded RNA genome. B-D) Negative controls where B) virus, C) DNA or D) $CaCl_2$ were replaced with water were also observed. Scale bar 10µm. E) Schematic representation of a baculovirus particle. Virus particles are rod-shaped, with a single transmembrane protein (GP64), beneath which lies a nucleocapsid which encloses the circular double-stranded DNA genome. F-H) Negative controls where F) virus, G) DNA or H) $CaCl_2$ were replaced with water were also observed.

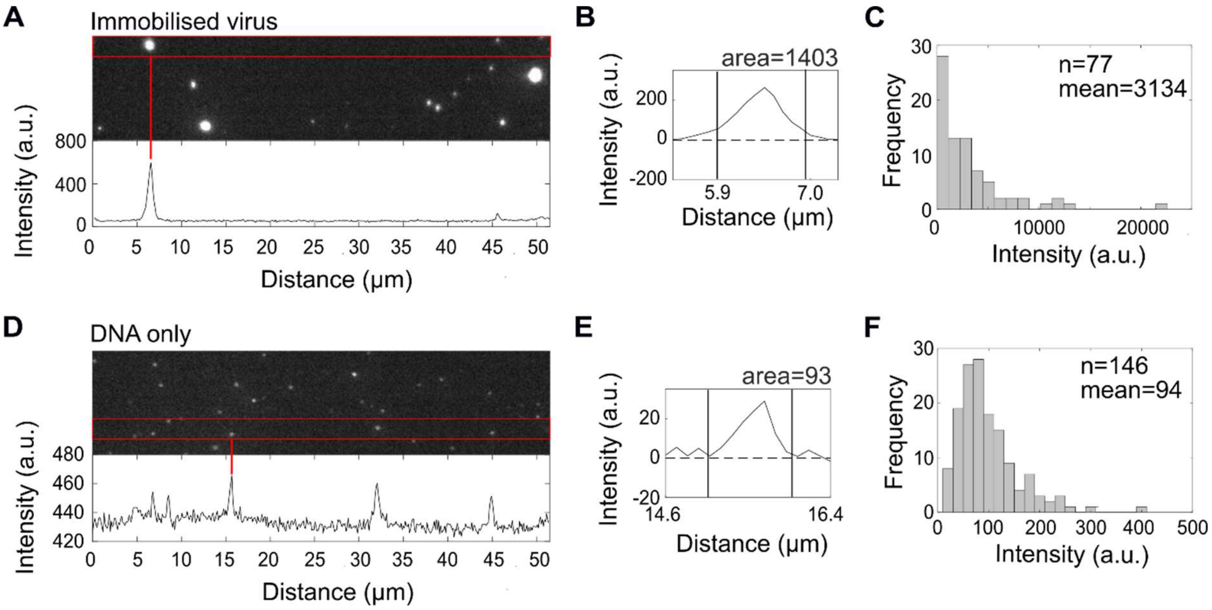


**Supplementary Figure 9. Fluorescently labelled viruses have multiple DNA molecules bound per virus particle.** A) A/Puerto Rico/8/1934 (H1N1) virus particles were incubated with 0.65M $CaCl_2$ and 1nM biotin-conjugated DNA labelled with Atto647N, before being immobilised on the surface of a pegylated glass slide via neutravidin. The upper panel shows a representative field-of-view of immobilised virus particles and the lower panel represents the intensity profile of the area indicated within the red box. B) Integration of the intensity peak corresponding to the virus particle highlighted in A). The dotted line indicates the local background threshold, which has been subtracted from the intensity profile, and the solid lines indicate the integration limits. C) Histogram of the integrated intensities of multiple virus particles. Number of virus particles (n) = 77 and mean intensity 3134 a.u. D-F) As for A-C but for DNA only. Number of virus particles (n) = 146 and mean intensity 94 a.u., resulting in an average number of 3134/94=33 DNA molecules per virus particle.

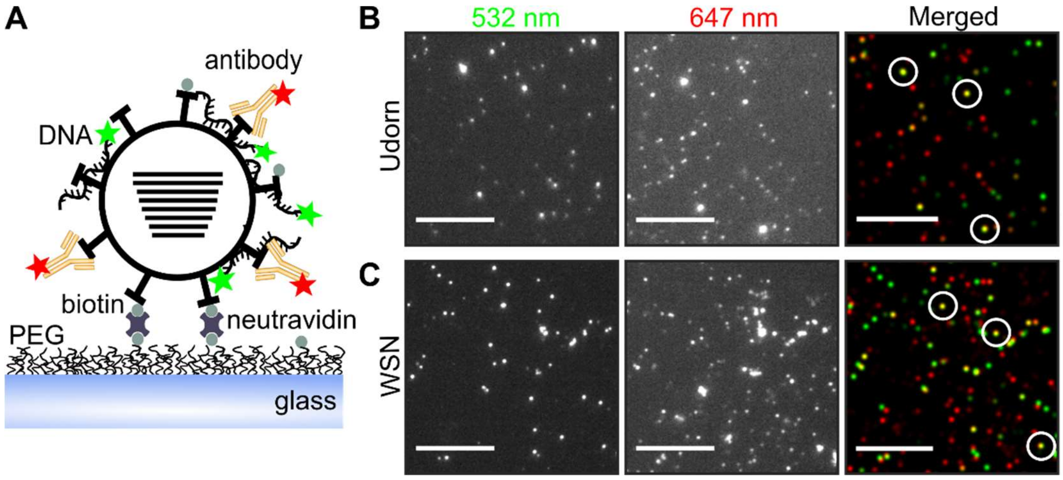


**Supplementary Figure 10. Calcium labelling of influenza viruses can be combined with specific antibody labelling.** A) Schematic representation of the assay, where green fluorescent DNAs were used to non-specifically label virus particles immobilized via biotin/neutravidin on the surface of a pegylated (PEG) glass slide. Immobilised viruses were then stained with red fluorescent antibodies specific to the virus strain. B) Representative field-of-view of A/Udorn/72 (H3N2) (Udorn) virus particles; calcium-labelling detected using a green 532 nm laser (left panel), antibody-staining detected using a red 647 nm laser (middle panel) and merged red and green localisations shown in the right panel. A/Udorn/72 virus at a final concentration of 12.5x$10^5$ PFU/mL was added to 0.65M $CaCl_2$ and 1nM Cy3B-labelled DNA before being immobilized. Viruses were stained with an anti-NA primary antibody and Alexa647-labelled secondary antibody before being observed on a wide-field microscope. White circles represent examples of merged localisations. Scale bar 10µm. C) Same as B), but with 41.2x$10^6$ PFU/mL A/WSN/33 (H1N1) (WSN) virus particles. Viruses were stained with an anti-NP primary antibody and Alexa647-labelled secondary antibody.

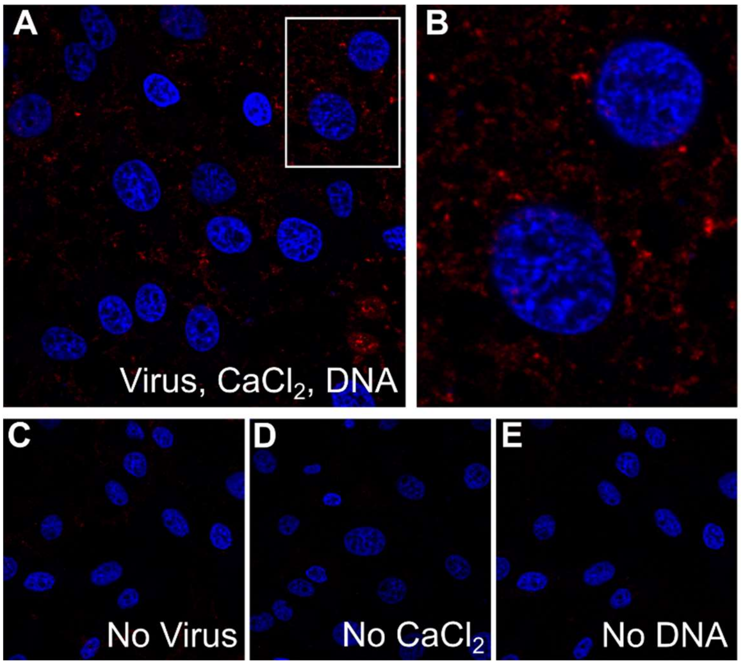


**Supplementary Figure 11. Fluorescently-labelled influenza viruses are able to adsorb to mammalian host cells.** A) A/Puerto Rico/8/1934 virus was added to 0.7M $CaCl_2$ and 1nM Atto647N-labelled DNA before being used to infect MDCK cells at a multiplicity of infection of ~$20x10^6$ PFU/mL. The cells were incubated for 1 hour at 37°C to allow viruses to adhere before being fixed and imaged. Cell nuclei were DAPI-stained and are coloured blue, cell cytoplasm occupies the regions between the nuclei. B) Zoomed in image of two cells (inset from A). Similar experiments where C) virus, D) $CaCl_2$ or E) fluorescently-labelled DNA were substituted with water were also carried out.

**Table 1: Sequences of DNA and RNA used in this study.**

| Name | Sequence (5' - 3') | Dye |
|---|---|---|
| DNA 1 | GAAATTGTTATCCGCTCTCACAATTCCACACATTATACGAGCCGAAGCATAAAGTGTCAAGCCX | X=T-Atto647N |
| DNA 2 | AGGCTTGACACTTTATGCTTCGGCXCGTATAATGTGTGGAATTGTGAGAGCGGATAACAATTTC | X=T-Cy3B |
| RNA | AGUAGAAACAAGGAGUUXUUUGAACAAACUACUU | X=U-Cy3 |
| 6mer | XCCACC | X=T-Atto647N |
| 9mer | XCCACCGTC | X=T-Atto647N |
| 12mer | XCCACCGTCGAT | X=T-Atto647N |
| 20mer | GGCAGTGAGCXCTACGCAAT | X=T-Cy3B |
| 45mer | CGCGAATGAGCTTACGTTATGTTACCTXATGTTACTTTAGATTTA | X=T-Atto647N |
| 64mer | Same as DNA 1 | X=T-Atto647N |
